# Manipulating virulence factor availability can have complex consequences for infections

**DOI:** 10.1101/062570

**Authors:** Michael Weigert, Adin Ross-Gillespie, Anne Leinweber, Gabriella Pessi, Sam P. Brown, Rolf Kümmerli

## Abstract

Given the rise of bacterial resistance against antibiotics, we urgently need alternative strategies to fight infections. Some propose we should disarm rather than kill bacteria, through targeted disruption of their virulence factors. It is assumed that this approach (i) induces weak selection for resistance because it should only minimally impact bacterial fitness, and (ii) is specific, only interfering with the virulence factor in question. Given that pathogenicity emerges from complex interactions between pathogens, hosts, and their environment, such assumptions may be unrealistic. To address this issue in a test case, we conducted experiments with the opportunistic human pathogen *Pseudomonas aeruginosa*, where we manipulated the availability of a virulence factor, the iron-scavenging pyoverdine, within the insect host *Galleria mellonella*. We observed that pyoverdine availability was not stringently predictive of virulence, and affected bacterial fitness in non-linear ways. We show that this complexity could partly arise because pyoverdine availability affects host responses and alters the expression of regulatorily linked virulence factors. Our results reveal that virulence-factor manipulation feeds back on pathogen and host behavior, which in turn affects virulence. Our findings highlight that realizing effective and evolutionarily robust anti-virulence therapies will ultimately require deeper engagement with the intrinsic complexity of host-pathogen systems.

## Introduction

The pervasive idea that virulence – the damage a host experiences during infection – follows more or less directly from pathogen load has shaped our view of infectious disease since the early days of germ theory (Bastian 1875; Pasteur 1880; Evans 1976; Anderson and May 1979; Frank 1996; Stearns and Koella 2008) and has underpinned our clinical quest to eradicate harmful microbes (Dagan et al. 2001; Allison et al. 2011; Russell 2011). However, advances over the years have revealed that the severity of an infectious disease depends on much more than just the sheer number of pathogens present: rather, it derives from complex interactions between the pathogen, its host, and the prevailing abiotic and biotic ecological conditions (Schmid-Hempel 2011; Bull and Lauring 2014; Méthot and Alizon 2014; de Lorenzo 2015). In other words, a microbe’s pathogenicity is not so much about what it is and how abundant it is, but what it does, when it does it, and to whom.

These insights have important consequences for antibacterial therapies that seek to control rather than eradicate infections (Vale et al. 2016). In particular, “antivirulence” approaches have been seen as promising alternatives to classic antibiotics (Cegelski et al. 2008; Rasko and Sperandio 2010; Allen et al. 2014; Vale et al. 2016). Such therapies seek to disarm rather than kill pathogens and do so by inhibiting the synthesis or the functioning of virulence factors (e.g. toxins, tissue-degrading enzymes, iron-scavenging siderophores, quorum sensing signals; Rahme et al. 1995; Miethke and Marahiel 2007; Nadal Jimenez et al. 2012; LaSarre and Federle 2013). The appeal of this strategy is that any effects on bacterial fitness should be relatively minor and therefore such treatments should induce only relatively weak selection for resistance (André and Godelle 2005; Pepper 2012). However, given the above-mentioned complexities intrinsic in infectious diseases, we can expect that in many cases a given antivirulence drug will have effects that extend beyond simply quenching the targeted virulence factor. We might have all sorts of unanticipated secondary effects on the behavior of the pathogen and its host. For example, the suppression of one virulence factor could pleiotropically affect the regulation of another virulence factor due to regulatory linkage at the genetic level (Herrera et al. 2007; Nadal Jimenez et al. 2012; Balasubramanian et al. 2013; GarÍia-Contreras et al. 2014). Furthermore, virulence factors often serve as cues for hosts to mount an immune response (Schmid-Hempel 2005; Park et al. 2014; Miyashita et al. 2015; Taszlow and Wojda 2015), so interfering with some virulence factor’s availability could indirectly modulate host responses.

In the light of this inherent complexity, it seems challenging to predict how a specific antivirulence therapy will likely affect bacterial load and treatment efficacy. If indeed the treatment causes secondary effects of the sort envisaged above, we might need to carefully reevaluate previous claims on the evolutionary robustness of such therapies. Complex interactions between pathogen and host factors could bring into play a multitude of different traits, all of which would be potential targets upon which natural selection could act on. Consequently, there could still be considerable selection for pathogen variants that are resistant to the treatment and/or become more virulent (Vale et al. 2014; Vale et al. 2016).

Here, we use the opportunistic human pathogen *Pseudomonas aeruginosa* as a test case to investigate the consequences of manipulating virulence factor availability. This bacterium relies on a number of virulence factors to establish infections in animals and humans, including immune-compromised cystic fibrosis patients (Rahme et al. 2000; Lyczak et al. 2002; Papaioannou et al. 2013). One particularly well-studied virulence factor is pyoverdine, a siderophore secreted into the local environment to scavenge iron from host tissue (Meyer et al. 1996; Harrison et al. 2006; Cornelis and Dingemans 2013). Pyoverdine is a multifunctional molecule. It can be shared as public good between cells for iron upake to stimulate growth and biofilm formation (Buckling et al. 2007; Banin et al. 2008). It is also used as a signaling molecule to control its own expression, and the synthesis of two additional virulence factors, exotoxin A and protease IV (Lamont et al. 2002). Additionally, it can act as a toxin by interfering with mitochondrial iron homeostasis (Kirienko et al. 2015). For all those reasons, pyoverdine has been identified as a suitable target for anti-virulence therapies (Kaneko et al. 2007; Banin et al. 2008; Visca et al. 2007; Lamont et al. 2002; Ross-Gillespie et al. 2014; Bonchi et al. 2014; Bonchi et al. 2015). In this study, we manipulated the availability of pyoverdine in the context of experimental infections of greater waxmoth larvae (*Galleria mellonella*). We investigated how interference with this virulence factor affects (i) bacterial growth within the host; (ii) the host’s response to infections; (iii) the pleiotropic regulatory links to other virulence factors; and (iv) how these factors combine and determine the overall level of virulence the host experiences. Building from previous work, we reduced the *in vivo* availability of pyoverdine by supplementing bacterial inocula with gallium, an iron mimic that inactivates pyoverdine molecules by binding irreversibly to them in place of iron (Kaneko et al. 2007; Ross-Gillespie et al. 2014). In addition, we also explored pathogen and host responses under conditions of increased pyoverdine availabilities. This allows us to test more generally how predictive virulence factor availability is for disease severity.

## Materials and methods

### Strains and media

Our experiments featured the clinical isolate *P. aeruginosa* PAO1 (ATCC 15692), a pyoverdine-defective knock-out strain derived from this wildtype (PAO1*ΔpvdD*), and three derivatives of these strains engineered via chromosomal insertion (*att*Tn7::ptac-*gfp, att*Tn7::ptac-*mcherry*) to constitutively express fluorescent proteins – i.e. PAO1-*gfp*, PAO1-*mcherry*, and PAO1Δ*pvdD-gfp*. Overnight cultures were grown in 8 ml Luria Bertani (LB) medium in 50 ml Falcon tubes, and incubated at 37°C, 200 rpm for 16-18 hours. For all experiments, we subsequently diluted the overnight-cultures in 0.8% NaCl saline solution. For *in vitro* assays, we used iron-limited CAA medium (per liter: 5 g casamino acids, 1.18 g K_2_HPO_4_^*^3H_2_O, 0.25 g MgSO_4_^*^7H_2_O, 100 μgml^−1^ human-apo transferrin, 20 mM NaHCO_3_, and 25 mM HEPES buffer). Human-apo transferrin in combination with NaHCO3 (as co-factor) is a strong iron chelator, which prevents non-siderophore-mediated iron uptake. All chemicals were purchased from Sigma-Aldrich, Switzerland. Pyoverdine was isolated using the protocol by Meyer *et al.* (1997).

### Manipulation of pyoverdine availability

In both our *in vitro* and *in vivo* assays we reduced and increased pyoverdine availability by supplementing bacterial inocula with, respectively, either gallium nitrate or purified pyoverdine. Gallium is an iron mimic that inactivates pyoverdine molecules by binding irreversibly to them in place of iron. It thereby lowers pyoverdine availability in a dose-dependent manner (Kaneko et al. 2007; Ross-Gillespie et al. 2014). The addition of pyoverdine immediately increases availability after inoculation, which has been shown to stimulate bacterial growth *in vitro* (Kümmerli and Brown 2010). For *in vitro* experiments, we varied gallium and pyoverdine concentrations from 5 to 250 μM. For *in vivo* experiments, we prepared inocula with 10-fold higher concentrations, since we assumed that upon injection into a host larva’s haemolymph (a total volume of approximately 100 μl; Harding et al. 2013) our infection inoculum (a 10 μl volume) would become diluted by a factor of approximately ten. Hereafter we report *in vivo* concentrations as estimated final concentrations, adjusted to reflect this assumed 10-fold dilution.

### *In vitro* growth and pyoverdine assays

To assess how our treatment regimes affects pyoverdine availability and bacterial growth, we performed in *in vitro* growth assays. Overnight LB cultures (PAO1 and PAO1Δ*pvd*D) were washed twice and standardized for optical density (OD = 2), then inoculated at 10^−3^ dilution to iron-limited CAA supplemented with either gallium nitrate (Ga(NO_3_)_3_; 5, 10, 20, 50 and 250 μM) or purified pyoverdine (same concentrations), to respectively reduce or enhance the availability of pyoverdine. All conditions were carried out in four-fold replication. Growth was tracked over 24 hours (37°C) in 200 μl cultures in 96-well plates (BD Falcon, Switzerland) using a Tecan Infinite M-200 plate reader (Tecan Group Ltd., Switzerland). We measured OD at 600 nm and pyoverdine-associated fluorescence (400 ex | 460 em), every 15 minutes following brief shaking of the plate (30s, 3.5 mm orbital displacement). Since gallium increases pyoverdine fluorescence, we corrected fluorescence values using a previously published calibration curve (Ross-Gillespie et al. 2014).

### *In vivo* growth assays

Infections were performed following protocols described in Ross-Gillespie et al. (2014). Briefly, final instar *Galleria mellonella* larvae, standardized for mass and general condition, were surface sterilized with 70% ethanol, inoculated between the posterior prolegs (Hamilton syringe; 26s gauge sterile needle), and then individually (randomly) distributed to the wells of 24-well plate for incubation at 37°C. *In vivo* bacterial growth was assayed as per Ross-Gillespie et al. (2014), using GFP fluorescence signal as a proxy for growth. For this reason, we infected larvae with bacterial strains harboring a constitutively expressed *gfp*-marker (i.e. PAO1-*gfp* or PAO1Δ*pvdD-gfp*). Inocula (10μl) contained ~25 colony forming units (CFU) of either PAO1-*gfp* supplemented with gallium (50 μM or 250 μM) or pyoverdine (50 μM or 250 μM), no pyoverdine, or the pyoverdine-defective PAO1Δ*pvdD-gfp* as a control treatment. A growth-negative control included the injection of saline solution. At 17 hours postinfection, larvae (24 per treatment) were processed to estimate their bacterial load. Approximately 50% of the larvae that had been infected with the wildtype strain were already dead at this time point. Larvae were individually flash-frozen in liquid nitrogen and then ground to fine powder using sterile micropestles. Powderized larva homogenates were resuspended in 1ml sterile H_2_O and centrifuged at 6300 RCF for 2 minutes. Thereafter, 200 μl of the water-soluble liquid phase of each sample was transferred to a 96-well plate and assayed for GFP-associated fluorescence using a Tecan Infinite M-200 plate-reader. To examine whether the bacterial load at 17 hours post-infection is representative of within-host growth dynamics, we repeated the experiment for a subset of treatments (untreated wildtype, wildtype with intermediate (50 μM) gallium or pyoverdine concentration, pyoverdine-deficient mutant, saline control). At 13, 15, 17, 20 hours, we processed randomly-selected larvae (24 per treatment) as described above and measured their bacterial load.

### *Ex vivo* growth assays

We investigated the potential influence of host effects on bacterial dynamics via *ex vivo* growth assays in haemolymph. In a first step, we primed *G. mellonella* larvae by inoculating them with bacterial wildtype cultures featuring manipulated levels of pyoverdine (by supplementing inoculum with either intermediate (50 μM) concentrations of gallium or pyoverdine. As controls, we primed larvae by infecting them with either the pyoverdine-deficient strain, pyoverdine alone, heat-killed wildtype bacteria, or the saline control. In a second step, we then measured bacterial growth in haemolymph extracted from these primed larvae. The priming inocula were administered as per the infection protocol described above. Inoculated larvae were distributed, in groups of 4, to petri dishes and incubated at 37°C. After 14 hours, the petri dishes were placed on ice for 15 min to anaesthetize the larvae prior to haemolymph extraction. A small incision was made in the posterior segment using a sterile scalpel, and haemolymph was drained with the aid of gentle pressure (Harding et al. 2013). From each sample, 25 μl of hemolymph was immediately stabilized with 15 μl of an ice-cold pH 6.5 cacodylate buffer (10 mM Na-C_2_H_7_AsO_2_ and 5 mM CaCh) and 15 μl of a saturated propylthiouracil solution to inhibit melanisation. Samples were then centrifuged (514 RCF, 2 min) to separate the liquid haemolymph fraction from any solid tissue contaminants, and 30 μl aliquots were transferred to individual wells of a 96-well plate, each containing 70 μl of saline solution. To kill the priming strains and any other bacteria that may have been present in the haemolymph as part of the natural larval microbiota, we added gentamycin to the haemolymph/buffer mixture to a final concentration of 20 μg/ml (a concentration known to kill susceptible *P. aeruginosa*; Choi et al. 2005). Subsequently, we inoculated wells with bacteria from an overnight culture (adjusted to an OD = 2 and subsequently diluted to 10^−4^) of a gentamycin-resistant PAO1-*mcherry* strain (this strain showed the same growth p*att*ern as the untagged wildtype strain). The plate was transferred to a Tecan Infinite M-200 plate reader for 24 hours incubation at 37°C. Every 15 minutes, we measured cell density (measured via the *mcherry*-associated fluorescence: 582 ex | 620 em; note: using optical density as a proxy for cell density is not reliable in this naturally turbid medium). These experiments allowed us to ascertain (a) whether bacterial growth in haemolymph is affected by a hosts’ history of prior infection, and (b) whether the availability of pyoverdine during priming predicts subsequent bacterial growth. Note that residual pyoverdine from the priming inocula was below detection limit after haemolymph extraction, and therefore should not influence later bacterial growth patterns.

### Molecular investigation of pyoverdine-mediated pleiotropy

Because pyoverdine is not only a virulence factor but also a signaling molecule, manipulating pyoverdine availability might also affect, via interaction with the PvdS iron-starvation sigma factor, the production of two additional virulence factors, exotoxin A and protease IV (Ochsner et al. 1996; Wilderman et al. 2001; Lamont et al. 2002). We used qPCR to explore whether our extrinsic manipulation of pyoverdine levels could change pyoverdine-mediated signaling and therefore pleiotropically affect expression of genes for virulence factor production (*pvdS*, *toxA* coding for exotoxin A, *prpL* (alternative name: *piv*) coding for protease IV, and*pvdA* coding for one of the pyoverdine synthesis enzymes). PAO1 cells were grown until early-and mid-exponential growth phases in 20 ml standard CAA (in a sterile 500 ml Erlenmeyer) containing either (i) no supplement, (ii) 10 μM Ga(NO_3_)_3_ (to reduce pyoverdine availability), (iii) 200 μM purified pyoverdine (to increase pyoverdine availability), or (iv) 100 μM FeSO_4_ (our negative control under which pyoverdine production should be completely switched off; Kümmerli et al. 2009). RNA was extracted using a modified hot acid phenol protocol and purified as in (Pessi et al. 2007; Pessi et al. 2013). Residual DNA in the sample was eliminated using RQ1 RNAse-Free DNAse I, and purification was performed using the RNeasy Mini Kit (Qiagen). Absence of DNA was verified by PCR using the primers specified in Table S1, and 40 cycles with the GoTaq Polymerase (Promega, Switzerland). RNA quality in the purified samples was then assessed using RNA Nano Chips (Agilent 2100 Bioanalyzer; RIN (RNA-Integrity Number) >7.6). First-strand cDNA synthesis with10 μg of total RNA from each sample was performed with M-MLV reverse Transcriptase RNase H Minus (Promega) and random primers (Promega). cDNA was subsequently purified with the MinElute PCR Purification Kit (Qiagen). The expression of *Pseudomonas aeruginosa* PAO1 genes PA2399 (*pvdD*), PA2426 (*pvdS*), PA1148 (*toxA*) and PA4175 (*prpL*) was analyzed with a Stratagene MX300P instrument (Agilent) using GoTaq qPCR Master Mix (Promega). All PCR-reactions were analyzed in triplicates with 3 cDNA dilutions (15, 7.5 and 3.25 ng) and 0.4 μM of individual primers in a total volume of 25 μl per reaction. Primers (Table S1) were designed using Primer3Plus (Untergasser et al. 2007) and subsequently verified by PCR. Fold-changes were calculated using the ΔΔCT method (Pfaffl 2001) using the primary sigma factor *rpoD* (PA0576) as housekeeping gene for data normalization.

### Virulence assays

Infections were performed as described above. Inocula (10 μl) contained ~25 CFU of *P. aeruginosa* from an overnight culture, resuspended in saline solution and supplemented with either gallium nitrate (5 μM, 50 μM, 250 μM), pyoverdine (10 μM, 50 μM, 250 μM) or neither. Controls included saline-only, gallium-only (50 μM and 250 μM) and pyoverdine-only inocula (50 μM and 250 μM), and also the PAO1 *ΔpvdD* strain, defective for pyoverdine production. The vitality of all larvae (i.e. spontaneous movement / response to tactile stimulation) was assessed hourly, starting at 10 hours post injection. Some of the larvae (n = 25, 3.14%) either started pupating while under observation or died prematurely during the first 10 hours post injection – presumably as a result of handling – and hence were excluded from further analyses.

### Statistical analysis

We used the functions from the ‘grofit’ R package to fit spline curves to the growth and pyoverdine production trajectories. From these fitted curves, we extracted growth parameters. In particular, we focused on growth integrals (areas under curves), which combine information from the lag phase, growth rate and yield. Growth integrals are particularly useful for non-logistic growth trajectories as observed throughout our experiments.

Survival curves were analysed by fitting parametric Weibull survival curves with the aid of functions from the ‘survival’ R package (Therneau and Grambsch 2000). From the fitted models, we extracted the hazard ratios and use those values to estimate the mortality risk of larvae within each treatment. To confirm the robustness of our analysis we also performed Cox proportional hazards regression, which yielded qualitatively similar results.

We used both parametric and non-parametric statistical models to test for treatment effects. Specifically, we used Kendall rank correlation analyses to test for associations between pyoverdine availability, growth, host response and virulence The data from our *in vivo* and *ex vivo* growth experiments did not meet the criteria of normally distributed residuals and the homogeneity of variances, which precluded the use of parametric statistical tests. For these analyses, we used the non-parametric Kruskal-Wallis test. All analyses were performed in R 3.3.0 (R Development Core Team 2015).

## Results

### Treatment effects on *in vitro* bacterial pyoverdine availability and growth

We first tested whether our treatment regime (i.e. adding gallium to quench pyoverdine or supplementing additional pyoverdine) indeed altered pyoverdine availability as intended. We found that our treatment regime had a positive linear effect on pyoverdine availability (Fig. 1A, Kendall’s correlation coefficient: τ = 0.75, *p* < 0.001, measured during the first eight hours of the growth period when pyoverdine is most needed to overcome iron limitation Kümmerli and Brown 2010). Moreover, we found that our manipulation of pyoverdine availability had a significant linear effect on bacterial growth (Fig. 1B, τ = 0.93, *p* < 0.001): adding gallium reduced growth, while pyoverdine supplementation accelerated growth relative to the unsupplemented wildtype. Taken together, our *in vitro* experiments show that our treatment scheme successfully manipulates pyoverdine availability and that pyoverdine is a growth promoter, essential for bacteria to thrive in iron-limited medium.

**Figure 1.**
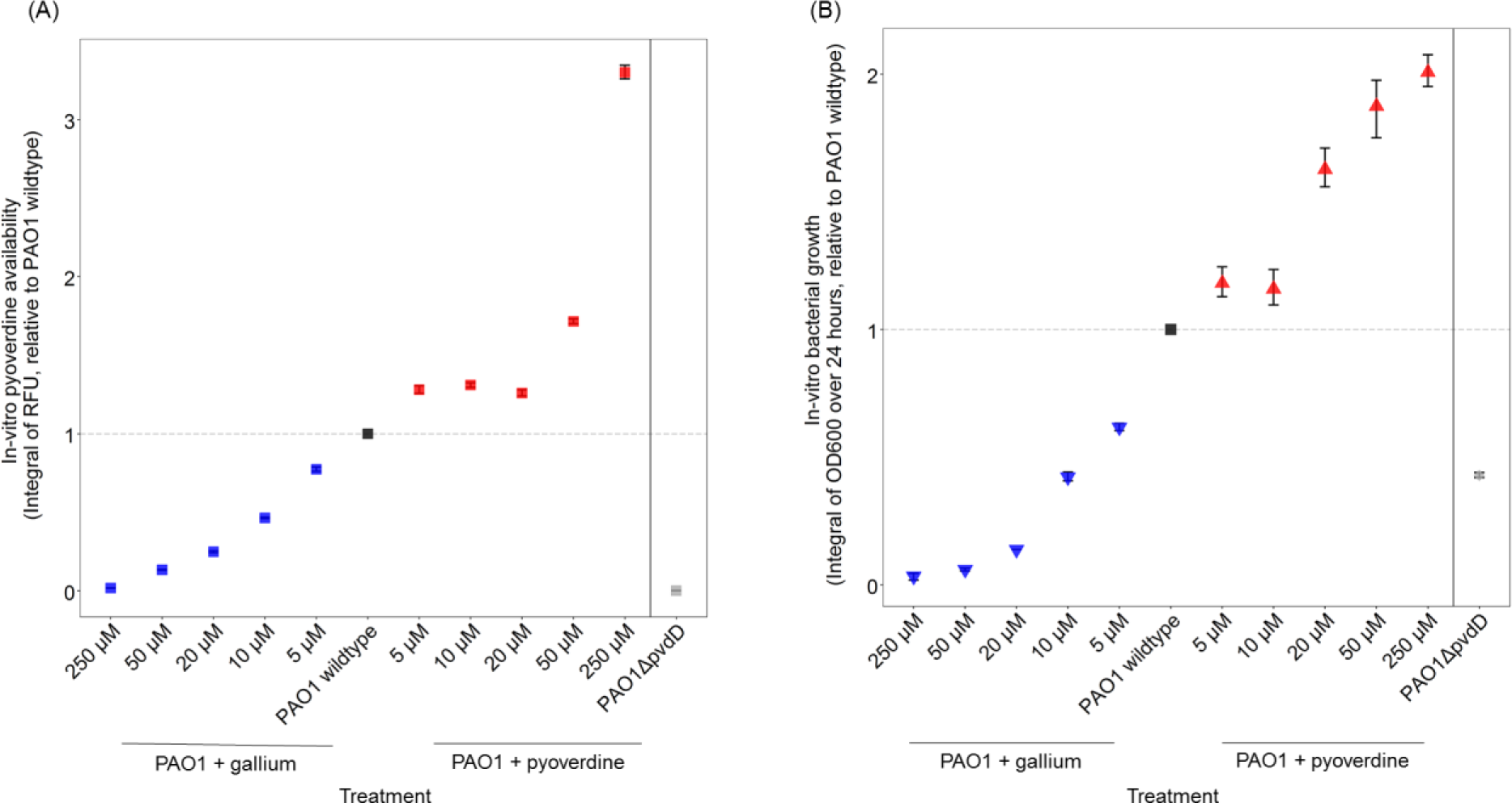
Our treatment regime significantly affected pyoverdine availability (A) and bacterial growth (B) in linear ways. To capture the dynamics of pyoverdine availability (first 8 hours) and growth (24 hours) in bacterial cultures, we used integrals (i.e. area under the curve) for analysis. Non-linear patterns in (A), e.g. for the 10 μM and 20 μM pyoverdine supplementations, can arise because bacteria plastically adjust their pyoverdine production level according to their need (kümmerli et al. 2009), such that *de novo* production and supplementation can balance each other out over time. Symbols and error bars represent mean estimates and 95% confidence intervals across four independent replicates.

### Non-linear effects of pyoverdine availability on *in vivo* bacterial growth

Pyoverdine availability also had significant effects on bacterial growth within the *G. mellonella* larvae (Kruskal-Wallis test for differences between treatments: *Χ*^2^ = 34.80, *p* < 0.001, Fig. 2), but the overall effect was not linear. Instead, bacterial load peaked in infections with the unsupplemented wildtype (i.e. at intermediate pyoverdine availability). Both the addition of gallium and pyoverdine significantly reduced bacterial growth compared to unsupplemented wildtype infections (for gallium 50 and 250 μM combined: *Χ*^2^ = 8.68, *p* = 0.013; for pyoverdine 50 and 250 μM combined: *Χ*^2^ = 6.66*, p* < 0.001). Bacterial growth also significantly peaked in infections with the unsupplemented wildtype when considering the entire growth trajectories and not only a single time point (Fig. S1), thereby confirming the above pattern.

**Figure 2.**
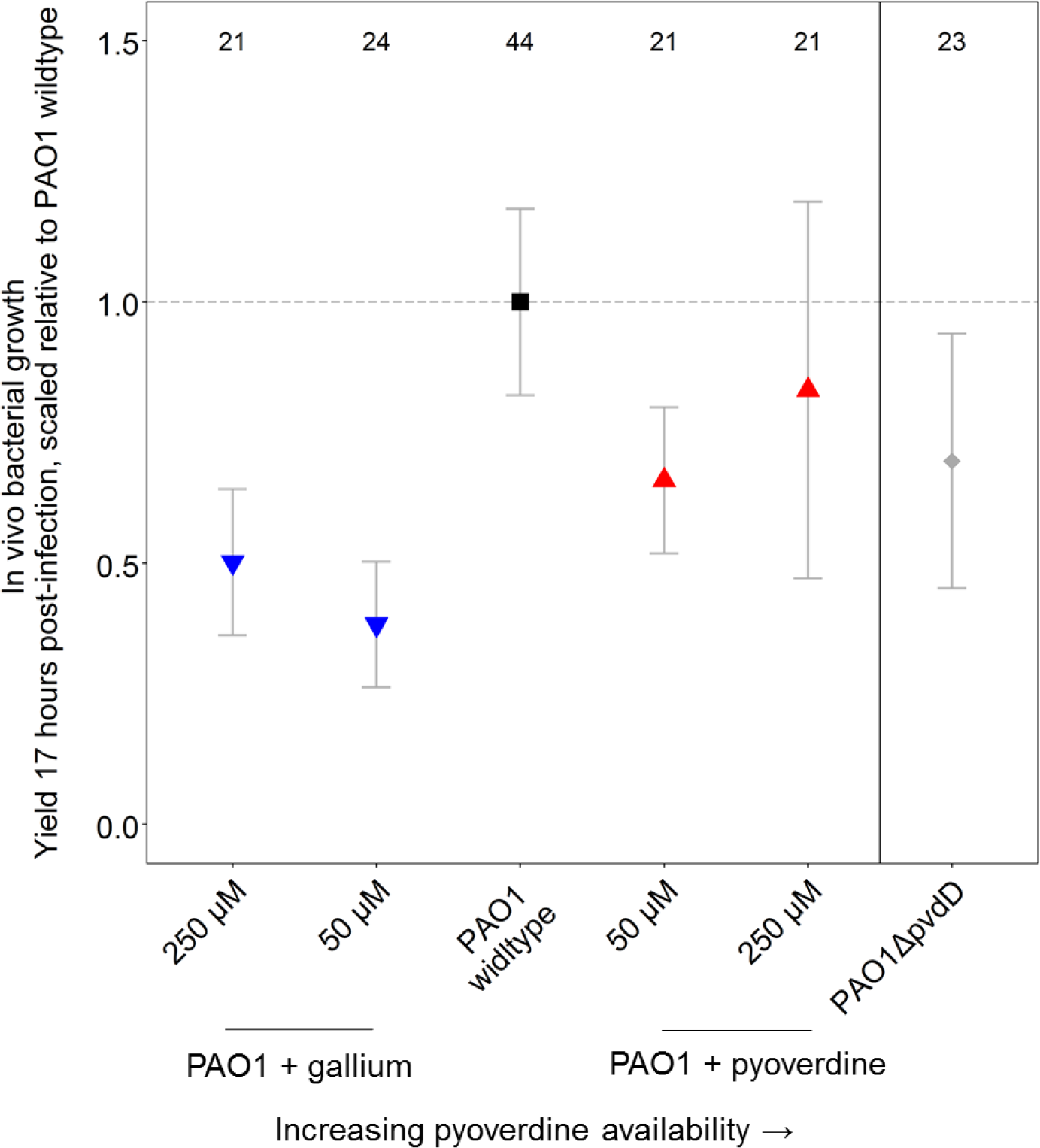
Pyoverdine availability has non-linear effects on *P. aeruginosa* growth in *G. mellonella* larvae. Bacterial load (measured 17 hours post-infection) peaked in infections with the unsupplemented wildtype, while the supplementation of both gallium (blue) and pyoverdine (red) resulted in a significant drop of bacterial load. Infections with a pyoverdine-deficient strain also resulted in a significant growth reduction compared to wildtype infections, indicating that pyoverdine is important for growth in this host. Symbols and error bars represent mean estimates and 95% confidence intervals, respectively. Numbers on top show sample size for each treatment.

One possible explanation for the absence of a linear relationship between pyoverdine availability and *in vivo* growth is that pyoverdine might not be required for bacteria to thrive within the host. However, two control experiments speak against this hypothesis. First, the growth of a pyoverdine-deficient knock out strains was significantly impaired in host infections compared to the wildtype (Kruskal-Wallis test: *Χ*^2^ = 7.54, *p* < 0.001, Fig. 2). Second, *ex vivo* growth of wildtype bacteria in extracted haemolymph demonstrated significant iron limitation and high pyoverdine production in this medium (Fig. S2). Altogether, these results indicate that pyoverdine is important for iron scavenging and growth within the larvae.

### Pyoverdine availability affects host responses

To investigate whether bacteria and/or pyoverdine availability triggers variation in host responses, we tracked growth of a wildtype strain *ex vivo* in haemolymph extracts from larvae previously primed under different conditions. *Ex vivo* bacterial growth in haemolymph indeed significantly differed depending on the infection history of the larvae (Fig. 3, Kruskal-Wallis test: *Χ*^2^ = 10.59, *p* = 0.014, including the pyoverdine manipulation regimes and the saline control). Specifically, bacteria showed significantly lower growth in haemolymph from wildtype-primed larvae than in haemolymph from saline-primed larvae (*Χ*^2^ = 4.11, *p* = 0.043). Furthermore, we found a significant negative association between the availability of pyoverdine in the priming inocula and the subsequent *ex vivo* bacterial growth (Fig. 3; Kendall’s τ = −0.21, *p* = 0.023). Control experiments revealed that a significant host response can be triggered by multiple stimuli: priming larvae with non-pyoverdine producing bacteria, pyoverdine alone, or heat-killed bacteria all resulted in a similarly increased response relative to the saline priming (Kruskal-Wallis test comparing pooled control treatments versus the saline treatment: *Χ*^2^ = 8.24, *p* = 0.004). Overall, our findings suggest that haemolymph primed with bacteria has a growth-inhibiting effect on *P. aeruginosa* and that this effect can vary plastically over time in response to pyoverdine availability.

**Figure 3.**
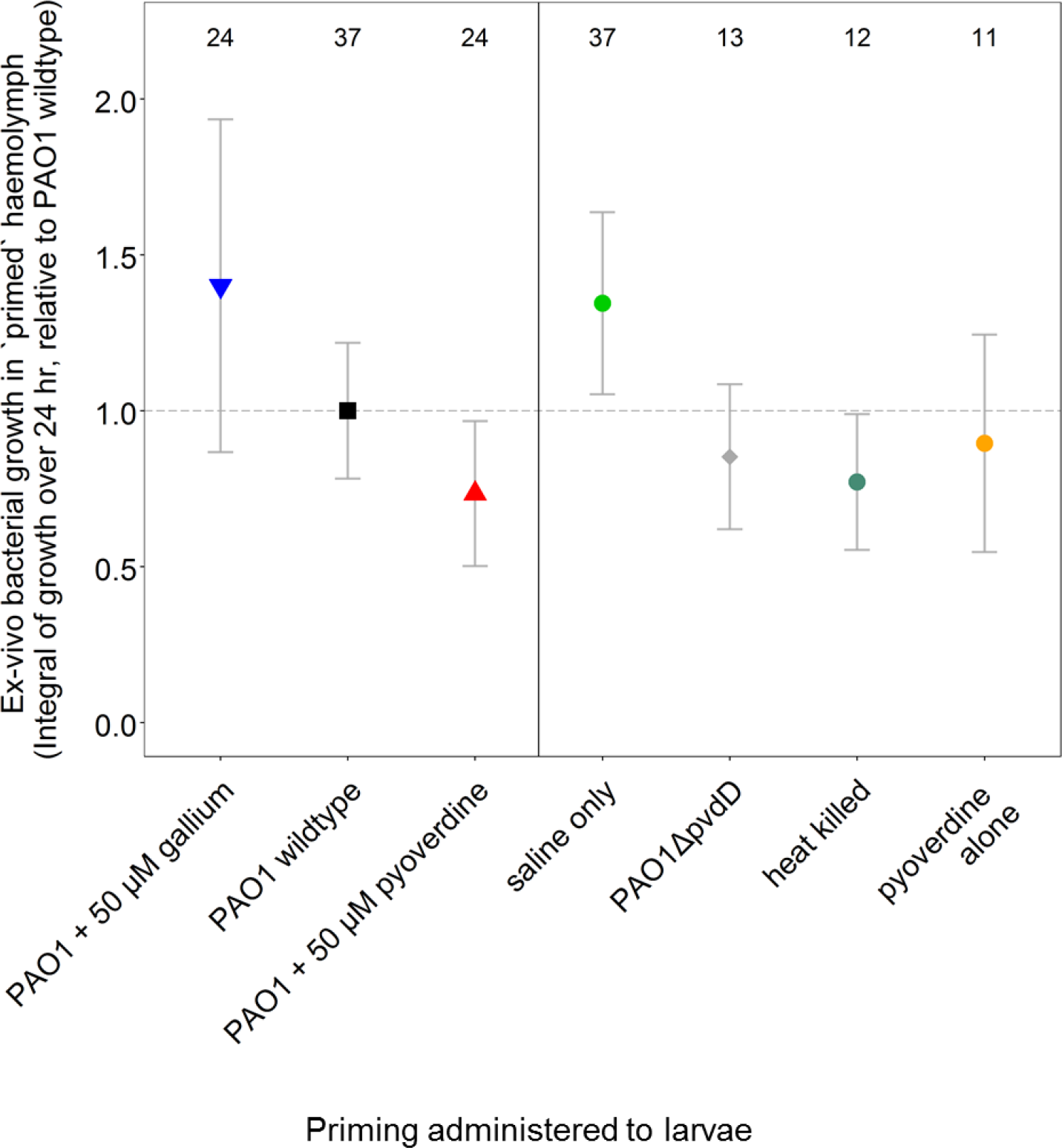
Growth of *P. aeruginosa* in haemolymph extracted from primed larvae demonstrates context-dependent host effects. Fourteen hours prior to haemolymph extraction, larvae were primed with wildtype bacteria, either alone or supplemented with gallium or pyoverdine. Control larvae were primed with saline alone, pyoverdine alone, heat-killed wildtype bacteria or a pyoverdine-deficient strain. Haemolymph extracts were gentamicin-treated to kill previously inoculated bacteria, and growth assays were then performed with a gentamicin-resistant wiltype strain. Compared to the saline-control, haemolymph primed with wildtype bacteria was significantly more refractory to subsequent bacterial growth, demonstrating a host response to infection. Moreover, we found a significant negative correlation between pyoverdine availability during the priming phase and the subsequent bacterial growth, indicating that pyoverdine is involved in triggering host responses. Symbols and error bars represent mean estimates and 95% confidence intervals, respectively. Numbers on top show sample size for each treatment.

### Pyoverdine availability affects the expression of other virulence factors

In addition to its function as a siderophore, pyoverdine is also a signaling molecule, which controls its own production and the synthesis of two other virulence factors, namely protease IV and exotoxin A (Lamont et al. 2002; Beare et al. 2002) (Fig. 4). It is therefore well conceivable that the experimental manipulation of pyoverdine availability also affects the expression of these other virulence factors. To test this hypothesis, we performed *in vitro* qPCR experiments, following the expression of the genes *pvdS*, *pvdA*, *prpL* and *toxA* across three levels of pyoverdine availabilities and two time points (early and mid-exponential phase). We examined these time points because pleiotropy relatively early in the growth cycle is likely to have the biggest effect on subsequent pathogen growth and virulence. The four genes code for the sigma factor PvdS (the main regulator of all three virulence factors), PvdA (enzyme involved in pyoverdine synthesis), protease IV, and exotoxin A (Fig. 4). Taking the unsupplemented wildtype bacteria growing in our standard iron-limited medium as a reference, we found that the addition of iron dramatically down-regulated the expression of all four genes (Table 1). This suggests that all three virulence factors (pyoverdine, protease IV and exotoxin A) are significantly expressed under the imposed iron-limited conditions (see also Ochsner et al. 2002). Next, we examined whether gene expression levels change as a function of pyoverdine availability. We found that pyoverdine manipulation either did not affect gene expression or resulted in the down-regulation of inter-linked genes (Table 1). Since there were no marked differences in gene expression profiles between the early and the mid-exponential growth phase, we pooled the data to identify the genes that were significantly down-regulated (Fig. 4). These analyses revealed that the addition of gallium (10 μM) slightly but significantly reduced the expression of *pvdS* (*t*_3_ = −10.55, *p* = 0.002) and*pvdA* (*t*_3_ = − 3.87, *p* = 0.031). The supplementation of pyoverdine (200 μM) significantly reduced the expression of *pvdA* (*t*_3_ = −17.95, *p* < 0.001) and *toxA* (*t*_3_ = −4.50, *p* = 0.020). Our results are promising from a therapeutic perspective, as they suggest that the manipulation of pyoverdine availability does not increase the expression of the interlinked virulence factors protease IV and exotoxin A, but rather has a neutral or even a negative effect on their expression.

**Figure 4.**
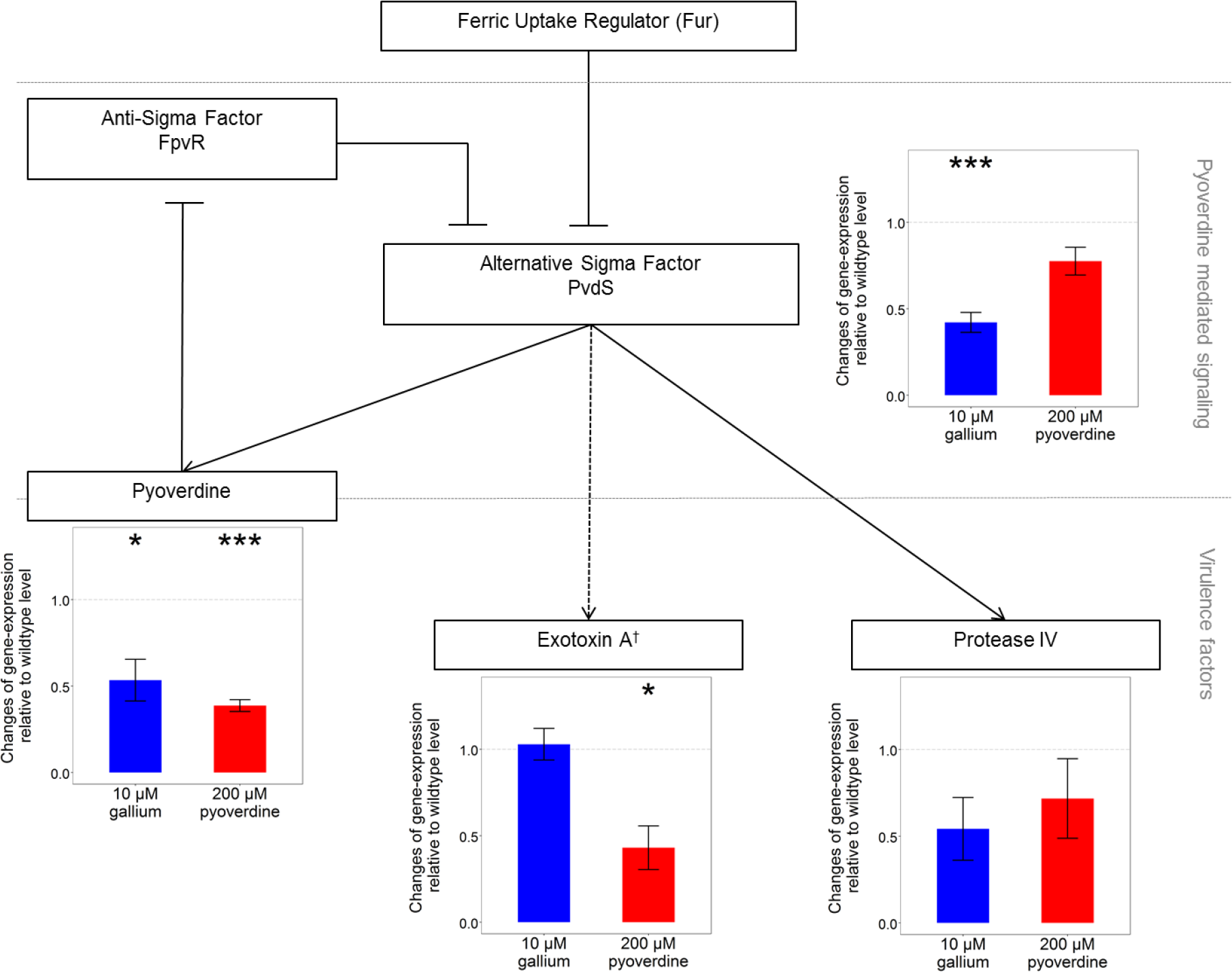
Manipulating pyoverdine availability has moderate effects on the pyoverdine-signaling network, and generally leads to the down-regulation of interlinked genes. Pyoverdine production is controlled by the alternative sigma factor PvdS that is itself negatively regulated by the ferric uptake regulator FUR, in response to intracellular iron levels. Pyoverdine modulates PvdS activity through a signaling cascade. Incoming iron-bound pyoverdine binds to its cognate receptor, thereby triggering the lysis of the membrane-bound anti-sigma factor FpvR, which binds and inhibits PvdS. In turn, membrane-released PvdS triggers increased transcription of pyoverdine synthesis genes, but also activates the expression of prpL (encoding the protease IV virulence factor), and toxR (coding for the ToxR regulator that then stimulates the expression of exotoxin A). Reduced pyoverdine availability (gallium supplementation, blue bars) moderately but significantly reduced *pvdS* and *pvdA* expression. Increasing pyoverdine availability (red bars) moderately but significantly reduced *pvdA* and *toxA* expression. Shown are mean values and standard errors across four replicates. Asterisks indicate significant gene expression changes relative to the unsupplemented wildtype (*p* < 0.05). † indirect regulation via *toxR*-regulator.

**Table 1.** Expression-fold changes for *P. aeruginosa* genes involved in pyoverdine-mediated signaling

| Growth Phase | Gene | Supplementation regime |  |  |  |  |  |  |  |
| --- | --- | --- | --- | --- | --- | --- | --- | --- | --- |
| | | | 100 μM FeSO<sub>4</sub> | | | 10 μM Ga(NO<sub>3</sub>)<sub>3</sub> | | 200 μM Pyoverdine | |
|  |  |  | mean | SE |  | mean | SE | mean | SE |
| Early-exponential | *pvdS* |  | 0.0003 | 0.0002 |  | 0.3256 | 0.0199 | 0.7237 | 0.1719 |
|  | *pvdA* |  | 0.0001 | 0.0000 |  | 0.3442 | 0.0353 | 0.3675 | 0.0159 |
|  | *toxA* |  | 0.0544 | 0.0098 |  | 0.9063 | 0.0161 | 0.2154 | 0.0200 |
|  | *prpL* |  | 0.2321 | 0.0375 |  | 0.7072 | 0.3661 | 1.1106 | 0.0821 |
| Mid-exponential | *pvdS* |  | 0.0003 | 0.0002 |  | 0.5167 | 0.0350 | 0.8273 | 0.0628 |
|  | *pvdA* |  | 0.0003 | 0.0002 |  | 0.7242 | 0.1163 | 0.4059 | 0.0775 |
|  | *toxA* |  | 0.0561 | 0.0092 |  | 1.1514 | 0.1412 | 0.6463 | 0.0519 |
|  | *prpL* |  | 0.0214 | 0.0046 |  | 0.3776 | 0.0896 | 0.3240 | 0.0143 |

Note: Expression fold-changes of *pvdS* (encoding the iron-starvation sigma factor PvdS), *pvdA* (coding for one of the pyoverdine synthesis enzymes), *toxA* (coding for exotoxin A), and *prpL* (encoding protease IV) are expressed relative to the unsupplemented PAO1 wildtype regime.

### Relationship between pyoverdine availability and virulence

Our results presented above (Figs. 2–4) show that the manipulation of pyoverdine availability has non-linear effects on bacterial load, triggers differential host responses, and has slight pleiotropic effects on the expression of other virulence factors. How do these factors now all combine within the host and determine the overall level of virulence associated with pyoverdine manipulation? Overall, our experimental infections of *G. mellonella* larvae revealed a significant positive association between pyoverdine availability and virulence (Fig. 5, Kendall’s τ =0.71, *p* = 0.030). Larvae died earlier in infections supplemented with pyoverdine, but survived longer when gallium was added instead. However, the trend was not altogether monotonic: moderately increased pyoverdine availability (10 μM) significantly decreased rather than increased the virulence risk (parametric survival regression assuming Weibull distribution: coefficient = 0.082 α 0.028, mean α SE, *z* = 2.89, *p* = 0.004). Such low-pyoverdine-supplementation infections showed virulence levels comparable to those of unsupplemented infections involving the pyoverdine-deficient PAO1*ΔpvdD* mutant (coefficient = 0.0055 α 0.028, *z* = 1.94, *p* = 0.846). The gallium-supplemented treatments were not ordered monotonically with respect to virulence, in that high (250 μM) gallium-supplemented infections were no less virulent than intermediate (50 μM) supplemented infections (coefficient = 0.043 α 0.037, *z* = 1.18, *p* = 0.238).

**Figure 5.**
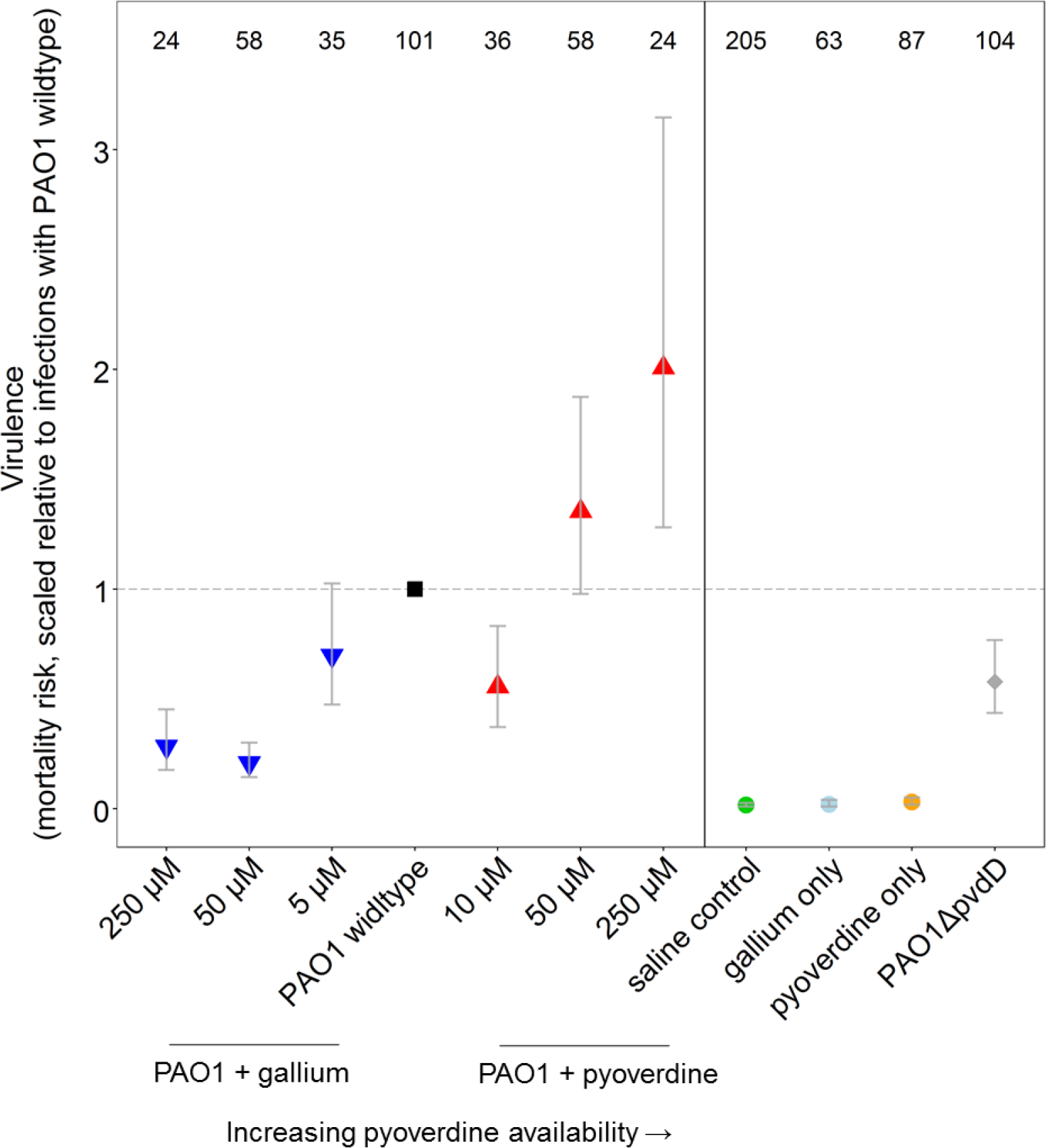
Relationship between pyoverdine availability and virulence, measured as mortality risk of larvae within each treatment. Overall, there is a positive correlation between pyoverdine availability and virulence with a notable exemption. Supplementing the infection with 10 μM pyoverdine reduced virulence comparable to a pyoverdine-deficient mutant. Symbols and error bars represent mean estimates and 95% confidence intervals, respectively. Numbers on top show sample size for each treatment.

## Discussion

Our results show that the manipulation of pyoverdine, an important virulence factor of the opportunistic human pathogen *P. aeruginosa*, affects bacterial load in infections of *G. mellonella* larvae in complex ways, triggers differential host responses, and influences the expression of other regulatorily-linked virulence factors (Figs. 2–4). Our findings have important consequences for recently proposed anti-virulence therapies, targeting pyoverdine-mediated iron uptake (Kaneko et al. 2007; Imperi et al. 2013; Ross-Gillespie et al. 2014; Bonchi et al. 2014; Bonchi et al. 2015), because complex interactions between bacterial load, host response and regulatory pleiotropy could result in unpredictable treatment outcomes (GarcÍa-Contreras et al. 2014). We examined this possibility for our system and found an overall positive relationship between pyoverdine availability and virulence, but also notable deviations from a monotonic pattern. For instance, the supplementation of low levels of pyoverdine significantly decreased rather than increased virulence, with this treatment reaching virulence levels comparable to infections with the pyoverdine-knockout strain (Fig. 5).

Some of the discovered complex non-linear associations between bacterial load, host response, pleiotropy and virulence warrant closer examination. For instance, why does increased pyoverdine availability (50 μM and 250 μM supplementation regimes) increase virulence despite the fact that these treatments reduce bacterial growth *in vitro* (Fig. 1B) and results, *in vitro* at least, in the down-regulation of the coupled virulence factor exotoxin A (Fig. 4)? One possible explanation is that high pyoverdine supplementation triggers an excessive host response, which is not only curbing bacterial growth, but is also damaging the host itself. For instance, *G. mellonella* produces the iron-chelator transferrin as part of its innate immune response (Han et al. 2004), a protein which actively counteracts the iron-scavenging activities of pathogens (Miethke and Marahiel 2007). Such a host response typically entails costs in terms of metabolic burden and autoimmune damage, and therefore must be appropriately calibrated (Day et al. 2007; Medzhitov et al. 2012). An overreaction from the host, perhaps in response to a high concentration of pyoverdine, could actually exacerbate, rather than reduce, virulence. Important to note is that although pyoverdine seems to induce a host response (Fig. 3) it is not toxic itself, as larvae infected with pyoverdine alone all remained healthy (Fig. 5).

Another complex association was that when increasing pyoverdine availability a little bit (10 μM) compared to the wildtype treatment, we observed a significant reduction of virulence (Fig. 5). This drop can potentially be explained by a host response too, but this time by a well-calibrated one, which primarily harms the pathogen while being beneficial for the host. If this explanation holds true, then the supplementation of moderate amounts of pyoverdine could represent a treatment that boosts host tolerance. Interestingly, treatments that increase host tolerance have, in addition to anti-virulence approaches, been proposed as alternative ways to combat infections (Medzhitov et al. 2012; Ayres and Schneider 2012; Vale et al. 2014; Vale et al. 2016).

Finally, we observed that infections with intermediate amounts of gallium (50 μM) were significantly less virulent than infections with the pyoverdine-deficient knock-out strain (Fig. 5). This suggests that this treatment has other effects, in addition to simply depriving siderophores from pathogens. One explanation would be that gallium has some general toxicity towards bacteria beyond its role in inhibiting iron uptake (Bonchi 2014). An alternative explanation, which is supported by our previous findings (Ross-Gillespie et al. 2014) but also the qPCR data (Fig. 4), is that intermediate gallium levels maintain pyoverdine synthesis, while high gallium levels completely stall the production. This steady production likely imposes a two-fold cost on bacteria: gallium does not only prevent pyoverdine-mediated iron uptake, but also induces continuous replacement of pyoverdine, which likely demands a high metabolic investment for very little reward (because pyoverdine is quenched by gallium once secreted). Given the ubiquity of linkages and feedback loops in the genetic architecture of bacteria (Nadal Jimenez et al. 2012; Dumas et al. 2013; GarcÍa-Contreras et al. 2014; Fazli et al. 2014), such features are likely important contributors to non-additive effects between pathogen behavior, fitness and virulence.

Given the complexities of host-pathogen relationships we have highlighted in this study, what could be the evolutionary consequences for anti-virulence therapies? The central tenet of this approach was that disarming rather than killing pathogens should induce weaker selection for resistance because it exerts only minimal effects on pathogen fitness (André and Godelle 2005; Pepper 2008; Rasko and Sperandio 2010; Stanton 2013). Our study demonstrates that anti-virulence approaches can in fact substantially modulate pathogen fitness (Fig. 2, see also Liu et al. 2008), which clearly offers natural selection the opportunity to favor pathogen variants that are partially or fully resistant to the treatment (see Maeda et al. 2012; Ross-Gillespie 2014; Allen 2014 for detailed discussion). One obvious evolutionary response of pathogens in response to virulence-factor quenching is to overproduce the virulence factor in question in order to outpace the quenching activity of the drug. Our results indicate that such an adaptation could affect the host in two different ways. If the increase in virulence factor production is substantial, this could lead to the evolution of a more virulent pathogen, which causes increased damage to the host in the absence of the treatment. Conversely, if the increase in virulence factor production is relatively small then it could positively stimulate host responses, which in turn could curb virulence. Evolutionary responses leading to increased virulence factor production would likely involve the modification of regulatory elements. As evidenced by our study, regulatory elements cannot only affect the expression of the targeted virulence factor, but also modify the expression of additional linked virulence factors in the same regulatory network (see Fig. 4). How exactly such regulatory linkage would alter global virulence factor expression profiles of a pathogen in a host, and how this feeds back on virulence cannot easily be foreseen, and might vary in response to the specific host stimuli present in an infection (Park et al. 2014). Taken together, our considerations show that we still have very limited understanding of the evolutionary consequences of antivirulence therapies. There is definitely a great need for controlled experimental evolution studies that measure selection pressures and adaptation patterns both at the proximate and ultimate level.

Given our dwindling supply of new antimicrobials, and the increasing prevalence of resistance to those we already have (Levy and Marshall 2004; Fischbach and Walsh 2009), creative approaches such as anti-virulence therapies are certainly required (Ross-Gillespie and Kümmerli 2014; Perron et al. 2015). To turn these ideas into effective and robust clinical therapies, however, we must delve deeper into the complexity of host-pathogen systems.

## Acknowledgements

We thank Martina Lardi for technical assistance, and three anonymous reviewers for constructive comments. RK was supported by the Swiss National Science Foundation (grant no. PP00P3-139164) and the Novartis foundation for medical-biological research (14B0457), MW was supported by DAAD; SPB was supported by HFSP (grant RGP0011/2014).

---

**Figure S1:**
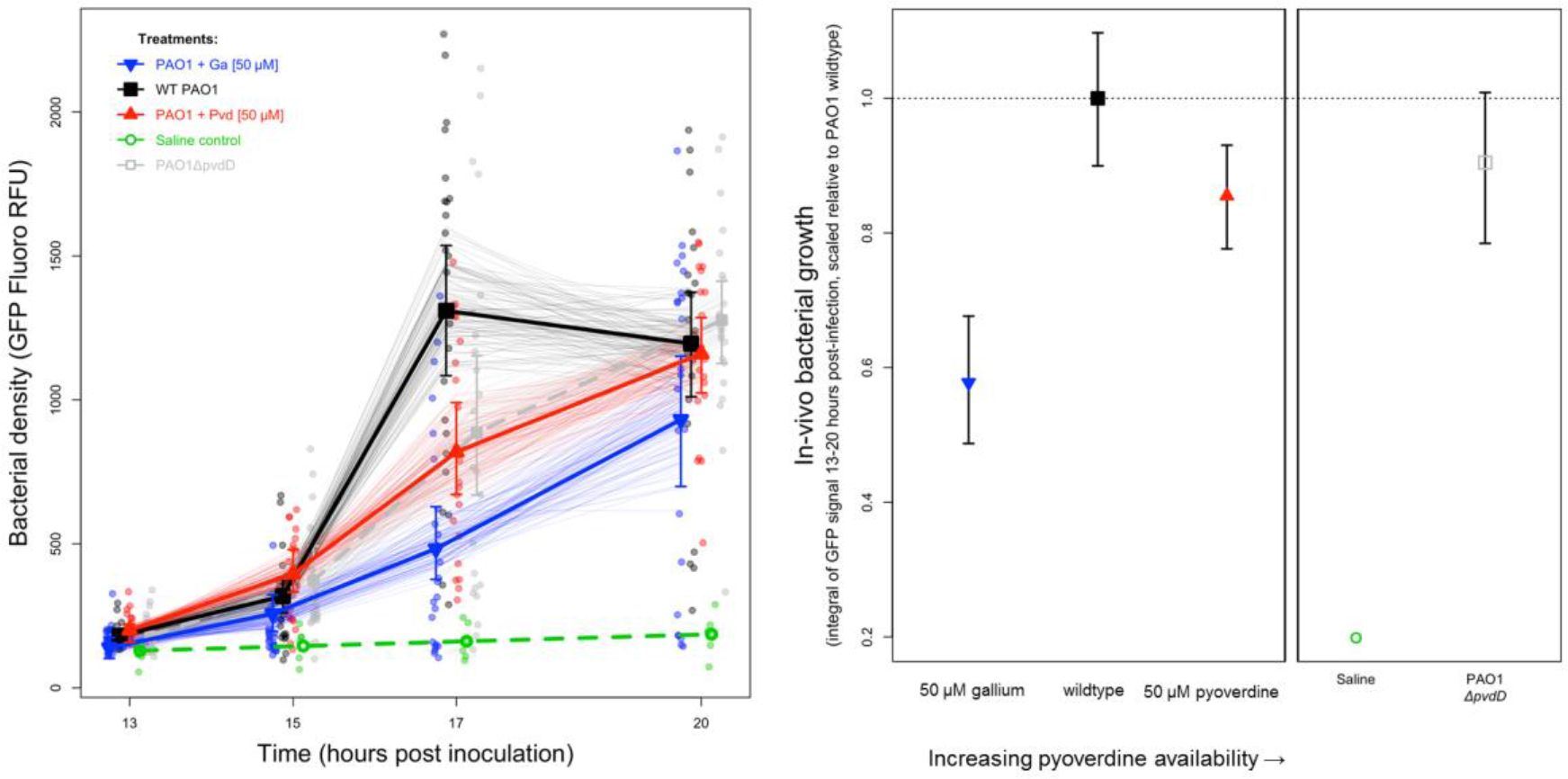
Growth trajectories of *P. aeruginosa* within *G. mellonella* larvae confirm that gallium and pyoverdine supplementation both significantly reduced bacterial growth compared to the unsupplemented wildtype (permutation test with 10,000 iterations: *p* =0.030). *In vivo* bacterial density was estimated from constitutively expressed GFP signal in host homogenates (points). This involved destructively sampling up to 96 larvae per treatment (~24 per time point; or n = 6 for the saline control). Because we were unable to track infections within an individual through time, we used bootstrap resampling of our observed data to generate replicated sets of estimated trajectories, a random sample of which are shown (faint lines). Symbols and error bars denote the medians, 2.5% and 97.5% quantiles from n=10,000 bootstrap-replicated datasets. We fitted splines to each trajectory and summarized the overall growth p*att*erns using areas-under-curves. These resulting distributions of these growth integrals are given in the final plot (medians with 2.5% and 97.5% quantiles).

**Figure S2:**
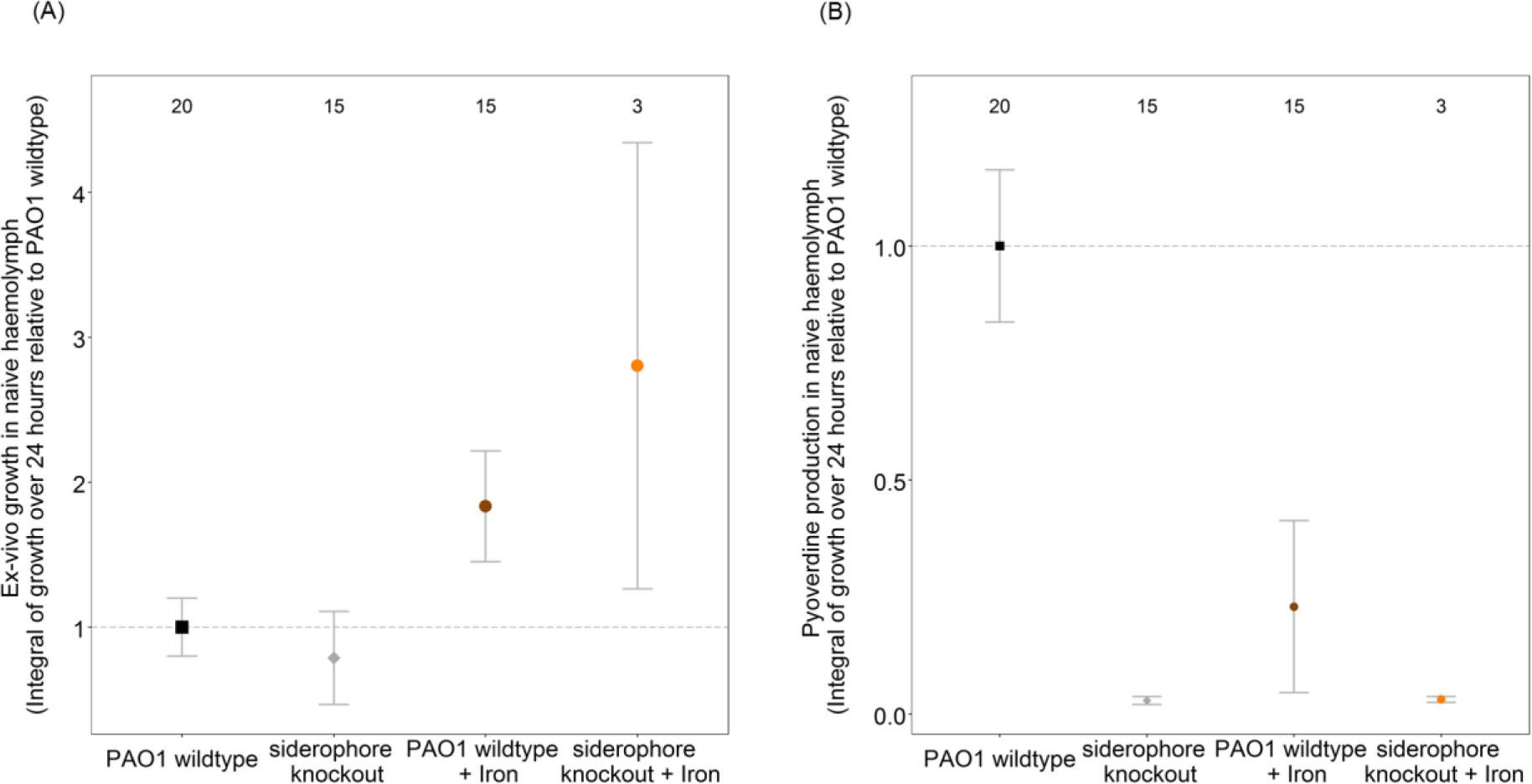
Growth (A) and pyoverdine production (B) of the wildtype strain and the pyoverdine-deficient mutant in naive haemolymph extracted from *G. mellonella* larvae. (A) The supplementation of 100 μM FeCl_3_ to the haemolymph significantly increased the growth of both the wildtype strain and the pyoverdine-deficient mutant, demonstrating that iron is a growth-limiting factor in the host environment. The observed pyoverdine-production profiles confirmed this assertion (B). Specifically, the wildtype strain produced high amounts of pyoverdine in the unsupplemented haemolymph, but reduced its investment to baseline level when iron was added to the haemolymph. Numbers on top show sample size of each treatment.

**S1 Table:**
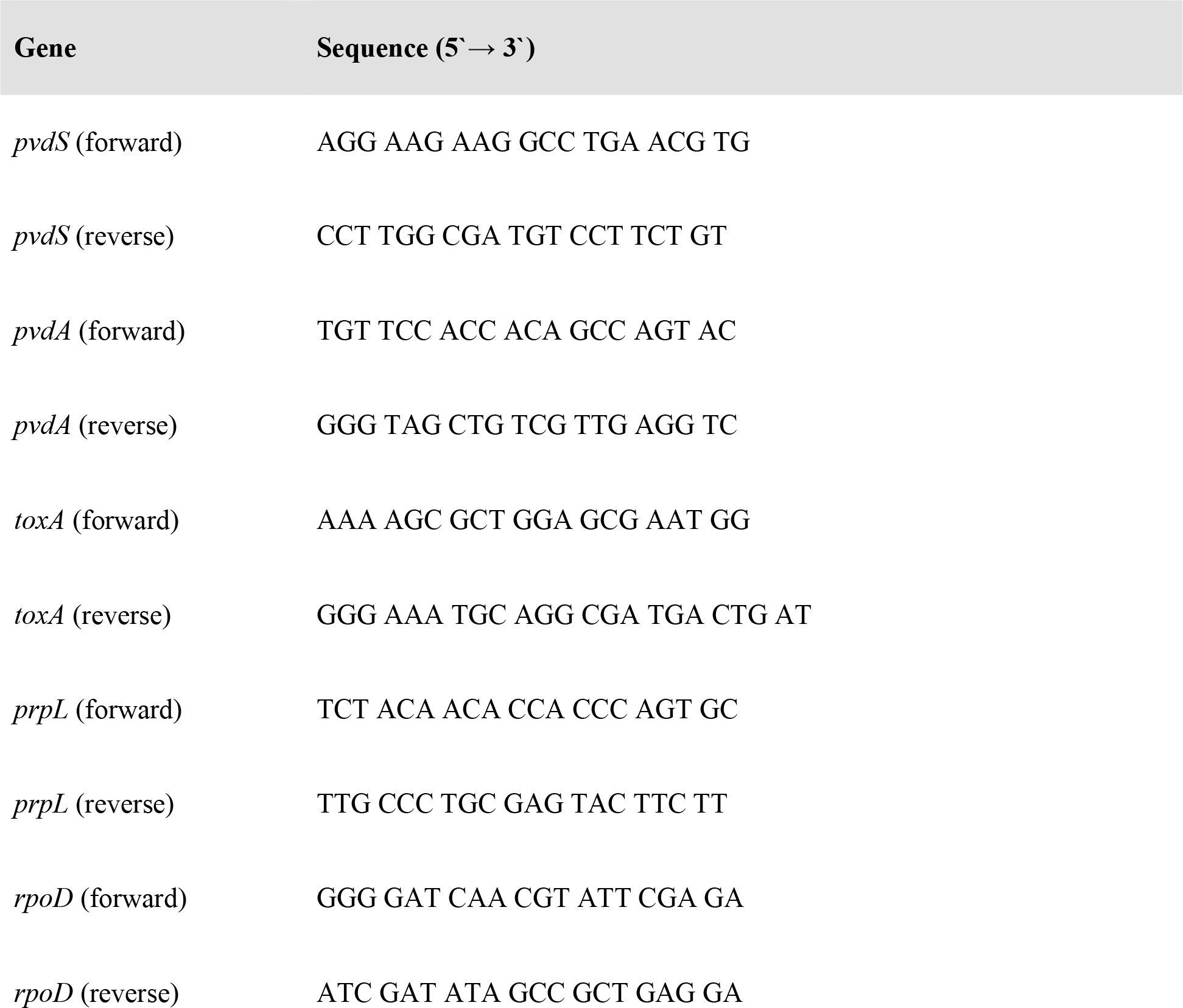
Genes and primers used for qPCR. We studied four genes involved in pyoverdine-mediated signaling, which are *pvdS* (encoding the iron-starvation sigma factor), *pvdA* (coding for an enzyme critical for pyoverdine synthesis), *toxA* (coding for exotoxin A), and *prpL* (encoding protease IV). Primers were designed based on sequences from the *Pseudomonas* genome database (http://www.pseudomonas.com) using the Primer3Plus platform aiming for 200 bp amplicons.

